# A Novel Metric for Quantifying the Sustainability of Phage-Mediated Bacterial Suppression

**DOI:** 10.64898/2026.08.14.744844

**Authors:** Tomoyoshi Kaneko, Daiki Tanaka, Shoichiro Koide, Yuna Tabata, Kazuhiko Miyanaga, Yasunori Tanji, Satoshi Tsuneda

## Abstract

The global spread of antimicrobial-resistant (AMR) bacteria represents one of the greatest threats to modern medicine, making the development of novel therapeutic strategies increasingly urgent. Phage therapy, which exploits bacteriophages (phages, viruses that specifically infect and kill bacteria) has regained attention as a therapeutic approach for multidrug-resistant infections. One critical determinant of treatment outcome is the capacity of phages to sustain bacterial growth suppression; however, no metric has previously existed to directly quantify the duration of effective lytic activity. Here, we propose the Sustainability Index (SusI), a novel metric that quantifies both the duration and extent of phage-mediated bacterial growth suppression, which is restricted to the primary lysis period from lysis initiation to resistance emergence. Evaluation of individual phages and two-phage cocktails against both laboratory and clinical isolates of *Escherichia coli* demonstrated that SusI provides information independent of the Virulence Index, which primarily reflects bactericidal activity during the initial phase of infection, and serves as a complementary metric to the Suppression Index, which may incorporate behavior beyond primary lysis. Cocktails composed of phages targeting different receptors specificities consistently exhibited higher SusI values, consistent with the notion that multifaceted selective pressure delays resistance emergence. Furthermore, in a mouse model of systemic infection established by intraperitoneal administration, cocktails with higher SusI values demonstrated superior therapeutic efficacy. These results confirm SusI as a practical metric for rational phage cocktail design. As phage therapy advances toward clinical implementation, standardized quantitative metrics such as SusI are expected to facilitate evidence-based selection of therapeutic phages across diverse pathogens and infection conditions.

**Importance:** The global spread of antimicrobial-resistant bacteria is making bacterial infections increasingly difficult to treat. Phage therapy, which uses bacteriophages (viruses that specifically infect bacteria), has re-emerged as a therapeutic alternative; however, reliable methods to determine in advance which phages will be therapeutically effective remain limited. Current evaluation metrics are well-suited for quantifying how rapidly phages kill bacteria but were not designed to directly measure how long lytic activity is sustained before resistant bacteria emerge. Here, we developed the Sustainability Index (SusI), a novel metric that specifically quantifies the duration of effective bacterial growth suppression. Evaluation of multiple phages and their combinations (cocktails) against both laboratory and clinical bacterial isolates demonstrated that SusI can distinguish phage combinations that existing metrics fail to differentiate. Moreover, in a mouse model of lethal bacterial infection, higher SusI values correlated with improved therapeutic outcomes. SusI has potential as a practical tool for selecting phages with greater likelihood of therapeutic success.

## INTRODUCTION

The emergence and rapid spread of antimicrobial resistance (AMR) represent one of the most serious challenges facing modern medicine. Recent estimates indicate that AMR infections cause more than 1.27 million deaths annually worldwide, with projections suggesting that this number could reach 10 million by 2050 if current trends continue (1). The World Health Organization has declared AMR one of the top ten global public health threats, underscoring the urgent need for therapeutic strategies that go beyond conventional antibiotics (2). Bacteriophage (phage) therapy, which exploits natural viruses that specifically target and kill bacteria, has attracted considerable attention as an alternative or adjunctive approach to antibiotic treatment (3, 4). Unlike broad-spectrum antibiotics, phages can target pathogenic bacteria with high specificity while preserving beneficial intestinal microbiota. Clinical successes, including compassionate use cases for multidrug-resistant infections, have demonstrated the therapeutic potential of phages and reignited scientific and clinical interest in this approach (5, 6).

A major challenge in phage therapy is the emergence of phage-resistant bacteria during treatment. Bacteria possess diverse defense mechanisms against phages, including receptor mutations, restriction-modification systems, and various other resistance strategies (7–11). Resistance acquisition frequently manifests as bacterial regrowth following initial suppression, which can lead to treatment failure and limit the clinical efficacy of phage therapy. To address this challenge, phage cocktails combining multiple phages have been proposed as a strategy to prevent resistance emergence (12). Recent studies have demonstrated that cocktails combining phages targeting different bacterial receptors can substantially reduce the probability of resistance emergence by imposing simultaneous selective pressure that requires the bacteria to acquire mutations in all targeted receptors (7, 13–15).

Currently, however, the metrics available for evaluating phage efficacy focus primarily on immediate bactericidal effects, and no established method exists for directly assessing the sustainability of lysis prior to resistance emergence. The Virulence Index (VI) and PhageScore quantify phage lytic activity based on bacterial reduction curves (16, 17). These metrics show a high correlation (r = 0.94), suggesting that they capture similar aspects of the phage–bacterium interaction during the acute phase of infection (18). While these metrics are useful for initial phage characterization, their application is limited when bacterial regrowth follows initial suppression, and may not adequately capture the sustained bacterial suppression and prevention of resistance that are essential for therapeutic success (19). The Suppression Index (SupI) is defined as the percentage of bacterial growth suppression over a fixed 30-hour measurement period (20); however, because SupI integrates data from the entire measurement period including behavior after resistance emergence, it cannot distinguish the persistence of primary lysis from subsequent resistant cell outgrowth. The temporal dynamics of phage–bacterium interactions are complex, encompassing multiple phases: initial phage adsorption and infection, bacterial lysis, resistance emergence, and subsequent bacterial regrowth. Existing evaluation metrics focus primarily on the early phases of this interaction, and no metric has previously existed to independently capture the sustainability of lysis prior to resistance emergence.

To address this limitation, we propose the Sustainability Index (SusI), a novel metric that quantifies both the duration and extent of bacterial growth suppression during the period from lysis initiation to resistance emergence. Here, we demonstrate the utility of SusI using a panel of phages with distinct receptor specificities: T1 and T5 (both targeting FhuA) (21, 22), T6 and ΦWec276 (both targeting Tsx) (23, 24), and ΦWec196 (targeting NfrA/B and lipopolysaccharide [LPS]) (25, 26). By comparing SusI with existing metrics (VI and SupI) across various phage combinations and multiplicities of infection (MOI), we establish SusI as a complementary tool for phage evaluation and cocktail design. Our findings suggest that SusI provides unique information about phage sustainability that current evaluation methods cannot capture, with the potential to improve the rational design of phage therapy regimens for clinical applications.

## MATERIALS AND METHODS

### Bacterial strains and phages

*Escherichia coli* MG1655 was used as the host strain and, unless otherwise specified, was cultured overnight in Luria-Bertani (LB) medium (BD Difco, Franklin Lakes, NJ, USA) at 37°C with shaking at 200 rpm. The clinical isolate of *E. coli* ESBL1064, obtained from a midstream urine sample at Gunma University Hospital, was also used as a host strain under the same culture conditions.

Phages T1, T5, T6, ΦWec196, and ΦWec276 were selected on the basis of their distinct receptor specificities. T1 and T5 both target FhuA (21, 22), T6 and ΦWec276 both target Tsx (23, 24), and ΦWec196 targets NfrA/B and lipopolysaccharide (LPS) (25, 26). Phages capable of infecting ESBL1064 (ΦWec430, ΦWec461, and ΦWec464) were isolated from urban sewage using the following procedure. Primary settling tank water (2 L each) was collected from four municipal wastewater treatment facilities and centrifuged (6,760 × g, 60 min, 4°C) to recover the supernatant. PEG 6000 (final concentration, 10 w/v%) and NaCl (final concentration, 4 w/v%) were added, and after overnight incubation at 4°C, phages were pelleted by centrifugation (12,000 × g, 90 min, 4°C). The pellet was resuspended in 10 mL SM buffer (50 mM Tris-HCl pH 7.5, 100 mM NaCl, 8 mM MgSO_4_·7H_2_O, 0.01% gelatin), and the aqueous supernatant collected after addition of an equal volume of chloroform, vortexing, and phase separation was stored at 4°C as concentrated wastewater stock. Phage isolation was performed by the double-layer agar method. The concentrated wastewater stock (100 µL) and an overnight culture of ESBL1064 (100 µL) were mixed into LB soft agar (0.5 w/v%) maintained at 50°C, overlaid onto LB agar plates (1.5 w/v%), and incubated overnight at 37°C. Individual plaques were purified through two rounds of streak culture on ESBL1064-seeded LB agar plates and stored. Additionally, phages from our laboratory’s phage library were screened for lytic activity against ESBL1064 by spot assay, and those demonstrating lytic activity were included in subsequent analyses.

### Antimicrobial susceptibility testing

The antimicrobial susceptibility of ESBL1064 was assessed by a microplate reader-based broth dilution method. Each antibiotic was added to LB medium at CLSI (Clinical and Laboratory Standards Institute) breakpoint concentrations (ampicillin, 16 µg/mL; amoxicillin, 16 µg/mL; cefalexin, 16 µg/mL; meropenem, 4 µg/mL), and turbidity changes were measured under the same conditions as the growth curve analysis described below.

### Phage propagation and stock preparation

Stocks of phages such as T1, T5, T6, ΦWec196, and ΦWec276 were prepared using MG1655 by a plate lysate method. An overnight culture of MG1655 (OD_660_ ≈ 2.0, 100 µL) and a high-titer phage suspension (100 µL) were mixed into 4 mL of molten LB soft agar (0.5 w/v%) maintained at 50°C. The mixture was immediately poured onto pre-warmed LB agar plates (1.5 w/v%), allowed to solidify at room temperature, and incubated overnight at 37°C until confluent lysis was observed.

To harvest phages from the plates, 5 mL of SM buffer was overlaid onto plates exhibiting confluent lysis and incubated at room temperature with orbital shaking at 500 rpm for 3 hours to allow diffusion of phages into the buffer. The buffer overlay was recovered by sterile pipette, transferred to a 15 mL conical tube, and centrifuged at 2,030 × g for 5 min at 4°C to pellet cell debris. The supernatant was collected, mixed with an equal volume of chloroform, vortexed for 30 sec, centrifuged at 2,030 × g for 5 min at room temperature, and the aqueous phase containing phages was transferred to a new tube.

PEG 6000 (FUJIFILM Wako Pure Chemical, catalog no. 167-22941) and NaCl (FUJIFILM Wako Pure Chemical, catalog no. 198-01675) were added to the phage-containing supernatant to final concentrations of 10 w/v% and 4 w/v%, respectively. After overnight incubation at 4°C, phages were pelleted by centrifugation at 9,000 × g for 30 min at 4°C. The supernatant was carefully decanted, and the phage pellet was gently resuspended in approximately 500 µL SM buffer by pipetting. An equal volume of chloroform was added, the mixture was vortexed and centrifuged at 2,030 × g for 5 min at room temperature, and the aqueous phase was recovered, supplemented with 100 µL chloroform, and stored at 4°C.

Stocks of phages such as ΦWec430, ΦWec461, and ΦWec464 were prepared using ESBL1064 by a liquid culture method. LB broth (30 mL) was dispensed into 50 mL centrifuge tubes, and overnight cultures of ESBL1064 (100 µL) and phage solution (100 µL) were added. Cultures were incubated overnight at 37°C with shaking at 200 rpm. The following day, cultures were centrifuged at 7,340 × g for 5 min at 4°C, and the supernatant was decanted into a 50 mL centrifuge tube containing 3 mL chloroform. After thorough vortexing, samples were centrifuged at 7,340 × g for 5 min at 4°C. The supernatant was recovered, and PEG 6000 and NaCl were added to final concentrations of 10 w/v% and 4 w/v%, respectively. After thorough mixing and overnight incubation at 4°C, samples were briefly vortexed and centrifuged at 7,340 × g for 30 min at 4°C. The supernatant was discarded, and the pellet was resuspended in 2–3 mL SM buffer. The resuspension was transferred to a 15 mL centrifuge tube containing 1 mL chloroform, vortexed, and centrifuged at 7,340 × g for 5 min at 4°C. The supernatant was collected and stored at 4°C.

Phage titers were determined by double-layer agar plaque assay. Phage stocks were serially diluted 10-fold in SM buffer, and 10 µL of each dilution was spotted onto plates. Plates were incubated overnight at 37°C, and plaques were counted at appropriate dilution steps. Phage titers were expressed as plaque-forming units per milliliter (PFU/mL). All phage stocks were diluted to appropriate concentrations in SM buffer to achieve the desired MOI in subsequent experiments.

### Generation of resistant mutants and estimation of receptor groups

To classify the target receptor groups for phages such as ΦWec430, ΦWec461, ΦWec462, ΦWec463, ΦWec464, ΦWec465, ΦWec466, and ΦWec467 upon infection against ESBL1064, resistant mutants of ESBL1064 were generated for each phage, receptor mutation-based resistance was confirmed by adsorption assay, and infection range assays were performed.

Resistant mutants were generated by two methods: liquid culture and double-layer agar. For the liquid culture method, a log-phase culture of ESBL1064 was diluted 100-fold, and a high-titer phage suspension (>10^8^ PFU/mL, MOI > 10, 100 µL) was added, followed by overnight incubation at 37°C with shaking at 200 rpm. Turbidity of the resulting culture was used to confirm resistant cell emergence, after which cultures were streaked onto LB agar. Two colonies per phage appearing the next day were selected, purified by streak culture at least twice on LB agar plates, grown in LB broth, and stored at −80°C. For the double-layer agar method, an overnight culture of ESBL1064 (100 µL) and a high-titer phage suspension (100 µL) were mixed into LB soft agar and overlaid onto LB agar plates, followed by overnight incubation at 37°C. Two resistant colonies per phage were selected and processed as described for the liquid culture method. In total, four resistant mutants per phage (two from each method) were obtained for each of the eight phages, yielding 32 resistant strains.

An overnight culture of each resistant strain (100 µL) was mixed with LB medium (890 µL), and the phage suspension used for resistance induction (>10^7^ PFU/mL, 10 µL) was added. Samples (10 µL) were collected at 0 and 10 min after phage addition and mixed with SM buffer containing chloroform (100 µL in 990 µL total), followed by centrifugation (15,000 rpm, ≥2 min, 4°C). The supernatant (100 µL) was mixed with an overnight culture of wild-type ESBL1064 (100 µL), plated in LB soft agar overlay, and plaque counts were determined after overnight incubation. The non-adsorption rate was calculated as the ratio of plaques at 10 min to those at 0 min; strains with an adsorption efficiency ≥95% (100 − non-adsorption rate) were designated receptor mutant strains.

Infection range assays of the receptor mutant strains were performed by spot assay. Each phage stock (>10^8^ PFU/mL) was serially diluted 10-fold (undiluted through 10^5^-fold dilution), and 1 µL of each dilution was spotted onto LB soft agar seeded with each receptor mutant strain. After overnight incubation at 37°C, the efficiency of plating (EOP) relative to wild-type ESBL1064 was calculated on a log scale for each receptor mutant strain. Strains with log EoP > −2.0 were considered susceptible, and the infection profile of each phage against the receptor mutant strains was compared to determine receptor group assignments. Additionally, spot assays were performed using the same procedure against an *Enterobacteriaceae* panel strain collection maintained in our laboratory comprising 45 *Enterobacteriaceae* strains (including laboratory and clinical *E. coli* isolates) to further support receptor group estimation by comparing host range patterns across phages.

### Growth curve analysis

Bacterial growth curves were measured using a pre-warmed (37°C) Epoch2 microplate reader (Agilent Technologies, Santa Clara, CA, USA). Overnight cultures were diluted 1:50 in fresh LB medium in I-shaped tubes and incubated at 37°C with shaking at 200 rpm until the optical density at 660 nm (OD_660_) reached approximately 0.5 (log phase), which typically required approximately 60 min.

For each experimental condition, 180 µL of diluted bacterial suspension (OD_660_ ≈ 0.1) was dispensed into wells of a 96-well clear flat-bottom microplate (Falcon®, Corning, NY, USA). Phage stocks were serially diluted in SM buffer to achieve the desired MOI (10^0^ to 10^-5^). Diluted phage suspension or SM buffer (for phage-free control conditions) (20 µL) was added to the corresponding wells, bringing the final volume to 200 µL per well.

Experiments using MG1655 were conducted in three independent biological replicates (overnight cultures prepared on different days), with 12 technical replicates per biological replicate (different wells on the same plate), for a total of 36 measurements per condition. Twelve wells per plate were used for phage-free control conditions to ensure reliable determination of control growth kinetics. For experiments testing phages and cocktails against ESBL1064, the number of biological and technical replicates varied depending on the phage and its isolation method; the specific numbers for each condition are provided in the figure legends.

Growth curves were monitored for 48 hours by measuring absorbance at 600 nm (OD_600_) at 15-min intervals, with continuous linear shaking (1,096 cpm, 1 mm amplitude) between measurements. The extended 48-hour measurement period was employed to capture the complete temporal dynamics, including bacterial growth, phage-mediated lysis, and potential resistance emergence. Detailed measurement conditions are provided in the Supplementary Materials.

### Calculation of bacterial growth indices

The Sustainability Index (SusI) was calculated by numerical integration using the following equation (Fig. 1):

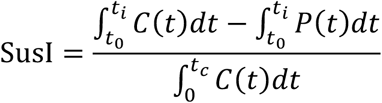

**Fig. 1.**
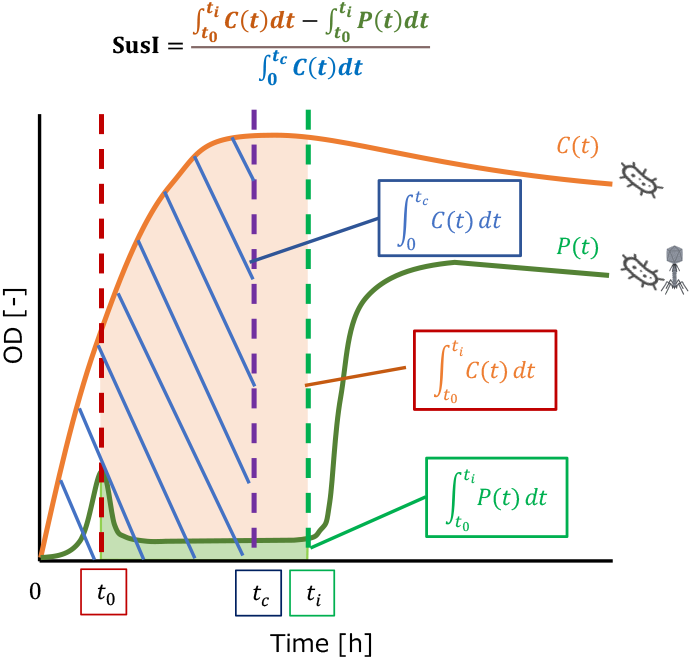
Schematic illustration of Sustainability Index (SusI) calculation. The orange curve represents the control bacterial growth curve *C*(*t*) and the green curve represents the phage-treated growth curve *P*(*t*). Key time points are indicated: *t*_0_ (lysis initiation time), *t_i_* (resistance emergence time), and *t_c_* (time to control stationary phase). The Sustainability Index is calculated as 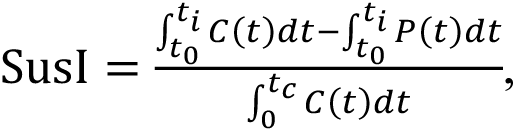 where the hatched area represents the denominator (the total area under the control curve from 0 to *t_c_*), and the numerator corresponds to the difference between the areas under *C*(*t*) and *P*(*t*) from *t*_0_ to *t_i_*. This index quantifies the sustainability of phage-mediated bacterial growth suppression relative to the duration of a normal growth cycle.

where *C*(*t*) represents the control growth curve without phage, *P*(*t*) represents the phage-treated growth curve, *t*_0_ is the lysis initiation time, *t_i_* is the resistance emergence time, and *t_c_* is the time at which the control culture reaches stationary phase.

The time points *t*_0_, *t_i_*, and *t_c_* were determined automatically from absorbance curves smoothed using a 3-point moving average: *t*_0_ and *t_i_* were identified by a sequential peak-detection algorithm, and *t_c_* was determined from the growth rate of the mean control curve using a threshold-based method. If ti was not detected within 24 hours, it was set to 24 hours, indicating that bacterial growth suppression was sustained throughout the measurement window. Full algorithm parameters are provided in the Supplementary Methods and Table S1.

The Virulence Index (VI) was calculated according to the method of Storms et al. (2020) (16), using the area under the bacterial reduction curve normalized by the area under the control growth curve during the initial phase of infection. The Suppression Index (SupI) was calculated following the methodology of Kim et al. (2024) (20), measuring the percentage of bacterial growth suppression over a 30-hour period. The time ratio (*t_i_* /*t_c_*) was calculated as a complementary metric to assess the duration of phage-mediated bacterial growth suppression relative to the normal growth cycle.

All indices were calculated using custom R scripts (R version 4.0.3; R Core Team, 2020) with the openxlsx, dplyr, ggplot2, and tidyr packages. The parameters used in these scripts are summarized in Table S1. Raw absorbance data output from the microplate reader were converted from seconds to hours prior to analysis.

### Mouse systemic infection model

All animal experiments were conducted with the approval of the Waseda University Animal Experiment Committee (approval number: A25-085(1)). In accordance with the 3R principles (Replacement, Reduction, and Refinement) for welfare of laboratory animals, the experimental protocol was carefully designed to minimize the number of animals required (Reduction) while maintaining sufficient power to detect robust biological trends. Female BALB/c AJcl mice (6 weeks old; CLEA Japan, Inc.) were used. To establish the infection model, serial concentrations of ESBL1064 prepared in phosphate-buffered saline (PBS; 10× concentrate from Nacalai Tesque, catalog no. 94222-61, diluted with MilliQ water and sterile-filtered through a 0.22 µm filter, pH 7.4) were administered by intraperitoneal injection (100 µL per mouse). A dose of 1 × 10^9^ CFU/mouse, which caused 100% mortality within 24 hours, was adopted as the infection dose. Administration of meropenem (30 mg/kg in 100 µL PBS) by tail vein injection simultaneously with infection resulted in 100% 7-day survival; this condition was therefore designated as the positive control (PC) for subsequent experiments.

For phage administration experiments, phage preparations in SM buffer (1 × 10^9^ PFU/mouse, MOI = 1, 100 µL) were administered by tail vein injection simultaneously with ESBL1064 infection (1 × 10^9^ CFU/mouse, 100 µL, intraperitoneally). Survival was recorded daily for 7 days and presented as Kaplan–Meier survival curves (n = 5/group). Two separate phage administration experiments were conducted. In the first experiment, three conditions with distinct combinations of VI and SusI (representative values at MOI = 1) were selected for comparison: T6 alone (low VI, low SusI), ΦWec461 alone (high VI, low SusI), and the T6+ΦWec461 cocktail (high VI, high SusI). In the second experiment, five conditions spanning a wide range of SusI values were selected to more systematically examine the relationship between SusI and survival: four cocktails (SusI = 1.272 ∼ 2.776) with consistently high VI (≈0.8) but differing SusI, together with the constituent single phage with the lowest SusI (ΦWec430 alone, VI = 0.501, SusI = 0.101), which also extended the range of VI examined. These were administered using the same protocol as the first experiment.

## Statistical analysis

Unless otherwise specified, statistical analyses for biological replicates were performed by first calculating the mean value for each biological replicate independently, then computing the standard error across the three biological replicates. Mouse Survival was recorded daily for 7 days and presented as Kaplan–Meier survival curves.

## Code availability

The R scripts used to calculate SusI, VI, SupI, and *t_i_*/*t_c_* are publicly available on GitHub at https://github.com/Tomoyoshi-Kaneko/susi-phage-analysis under the MIT License and are archived on Zenodo (https://doi.org/10.5281/zenodo.21890229).

## RESULTS

### Growth curve analysis and automated detection of time points in *E. coli* MG1655

*E. coli* MG1655 was co-cultured with five phages (T1, T5, T6, ΦWec196, and ΦWec276) individually or with two-phage cocktails (10 combinations), and turbidity curves were acquired over 24 hours in a 96-well plate format (Fig. 2, Fig. S1). The time to stationary phase of control cultures (*t_c_*) was detected in the range of 8–12 hours across all experiments (Fig. 2, Fig. S1, Fig. S2). All phages exhibited MOI-dependent lytic activity, with a general trend toward earlier lysis initiation at higher MOIs.

**Fig. 2.**
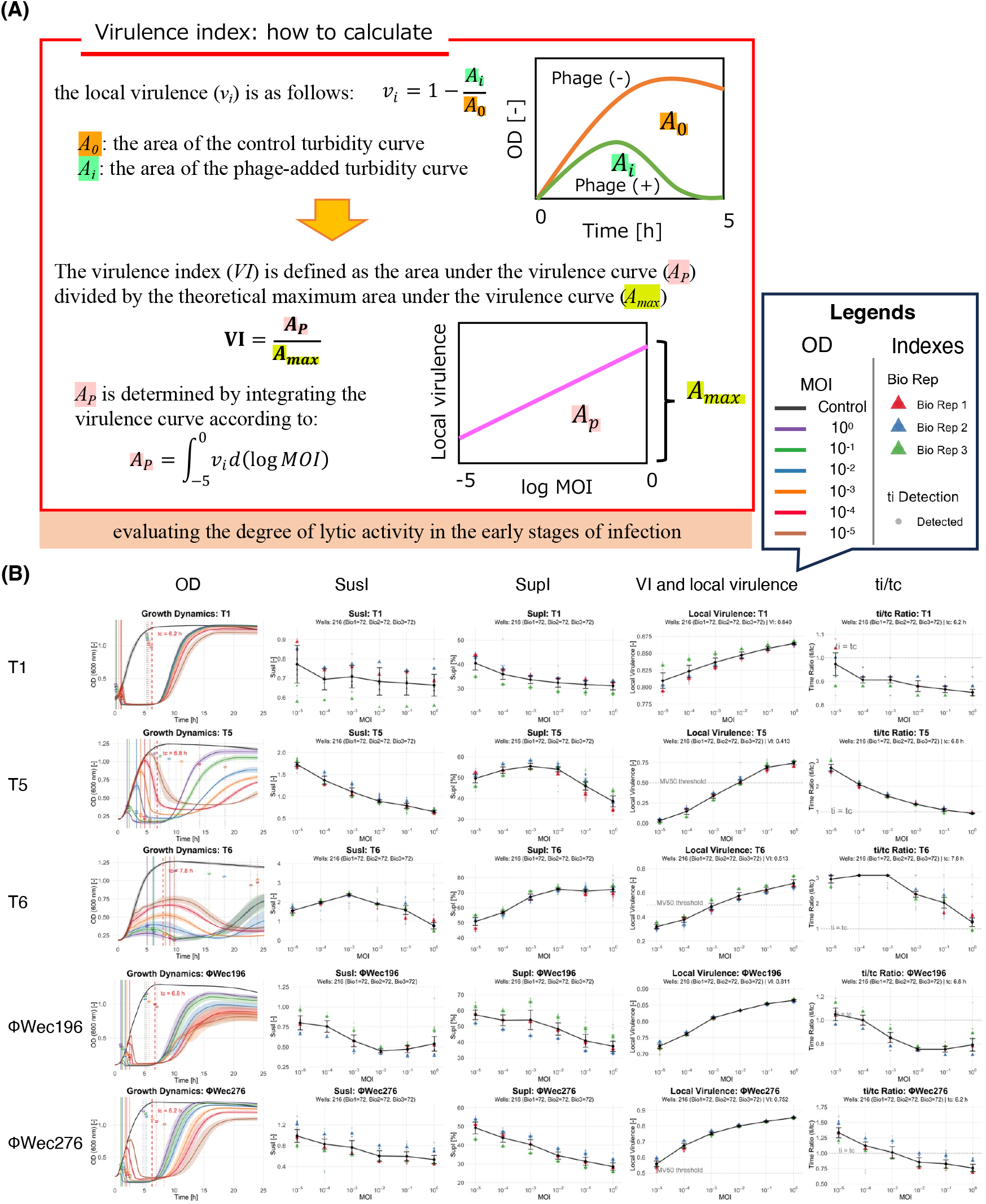
Growth curves and phage indices for single phages against *E. coli* MG1655. (A) Calculation of local virulence and the Virulence Index (VI), following the method of Storms et al. (2020) (16). Local virulence (*v_i_*) at a given MOI is defined as *v_i_* = 1 − (A_i_/A_0_), where A_0_ and A_i_ are the areas under the turbidity curve of the phage-free control and phage-treated culture, respectively. VI is then calculated as the area under the local virulence curve (A_p_), obtained by integrating vi over log (MOI) from −5 to 0, normalized by the theoretical maximum area (A_max_): VI = A_p_/A_max_. In panel (B), the MV50 threshold indicates the MOI at which local virulence reaches 0.5, analogous to an ID50; a lower MV50 indicates that a smaller amount of phage is sufficient to achieve substantial killing, i.e., greater potency. (B) Each row shows, from left to right, turbidity curves (OD_660_), SusI, SupI, Local Virulence (VI), and *t_i_*/*t_c_*. Turbidity curves are shown as mean ± standard error for each MOI condition (control and 10^0^ to 10^5^). In each index plot, semi-transparent circles (●) indicate values calculated from individual well turbidity curves, and triangles (▴) indicate values calculated from the mean curve of technical replicates within each biological replicate; both are color-coded by biological replicate (first: red, second: blue, third: green). Black diamonds (◆) and solid lines indicate values calculated from the mean turbidity curve across all biological replicates. Error bars represent standard error across three biological replicates. Phages analyzed are T1, T5, T6, ΦWec196, and ΦWec276.

### Comparison of indices for phages infecting MG1655

SusI, VI, SupI, and *t_i_*/*t_c_* were calculated and compared for the five single phages and ten two-phage cocktails (Fig. 2, Fig. S1). SusI quantifies the duration and depth of primary lysis by integrating the difference in area under the growth curve between the phage-treated and control groups from lysis initiation (*t*_0_) to resistance emergence (*t_i_*), normalized by the total area under the control curve to stationary phase (*t_c_*) (Fig. 1; see Materials and Methods for calculation details).

For single phages, SusI values at MOI = 1 fell within a range of 0.5–0.7 for T1 (0.664 ± 0.057), T5 (0.656 ± 0.030), ΦWec196 (0.539 ± 0.092), and ΦWec276 (0.535 ± 0.078), while T6 showed a slightly higher value (0.766 ± 0.163). Among cocktails, the same-receptor cocktail T1+T5 (0.856 ± 0.055) showed SusI values comparable to those of single phages, whereas different-receptor cocktails exhibited markedly elevated SusI values, with T1+ΦWec276 (2.059 ± 0.083), T1+ΦWec196 (1.614 ± 0.180), and T6+ΦWec196 (1.880 ± 0.465) showing particularly high values (Table 1).

**Table 1.** Target receptors of phages used against *E. coli* MG1655.

| Phage | Target receptor | Receptor function |
| --- | --- | --- |
| T1 | FhuA | Outer membrane ferrichrome-iron transporter (siderophore receptor) |
| T5 | FhuA | Outer membrane ferrichrome-iron transporter (siderophore receptor) |
| T6 | Tsx | Outer membrane nucleoside-specific channel |
| ΦWec276 | Tsx | Outer membrane nucleoside-specific channel |
| ΦWec196 | NfrA/B and LPS | Outer membrane protein complex (also serves as the receptor for phage N4) and lipopolysaccharide core |

VI was highest for T1 (0.840 ± 0.005) and ΦWec196 (0.811 ± 0.012) and lowest for T5 (0.413 ± 0.065). The VI values of the cocktails were similar to those of the constituent single phages, with no marked change upon cocktail formation. This absence of a clear difference in VI between single phages and cocktails stood in notable contrast to the behavior of SusI.

SupI for single phages was markedly higher for T6 (72.1 ± 1.6%) compared to the other four phages (28–40%). The SupI of multiple different-receptor cocktail combinations exceeded 80%, with T1+ΦWec196 (85.8 ± 2.2%) and T1+ΦWec276 (85.2 ± 1.1%) showing the highest values. The SupI of the same-receptor cocktail T1+T5 (33.6 ± 1.5%) remained comparable to single-phage values, a trend analogous to that observed for SusI. For T6 alone, *t_i_*/*t_c_* (1.258 ± 0.171) markedly exceeded 1, indicating sustained bacterial growth suppression throughout the observation period. All other single phages showed *t_i_*/*t_c_* < 1, indicating that resistant bacteria emerged before the control reached stationary phase. Different-receptor cocktails tended to show elevated *t_i_*/*t_c_* values, mirroring the trends observed for SusI.

### Correlation analysis between SusI and existing indices for phages infecting MG1655

The correlation coefficient between SusI and VI was 0.184, indicating no correlation between these two metrics (Fig. 3A). The correlation coefficient between SusI and SupI was 0.828, indicating moderate correlation, although multiple conditions deviated from the regression line (Fig. 3B). T6 alone showed relatively low SusI (≈0.77) compared to its SupI level (≈72%), and T1+T6 fell below the regression line with SusI ≈1.07 versus SupI ≈78%. Growth curve analysis revealed that T6 alone exhibits delayed lysis initiation compared to other phages, resulting in a smaller difference from the control during the early measurement period, and that T1+T6 undergoes secondary lysis — a re-decline in turbidity following resistance emergence (Fig. S1). The correlation coefficient between SusI and *t_i_*/*t_c_* was 0.979, indicating a strong correlation (Fig. 3C), though T6 alone represented an exception. Despite a relatively high *t_i_*/*t_c_* (1.258), T6 showed a low SusI (0.766) that fell below the regression line. Growth curve analysis confirmed that T6 maintained a higher turbidity level during the lysis period compared to other phages (Fig. 2).

**Fig. 3.**
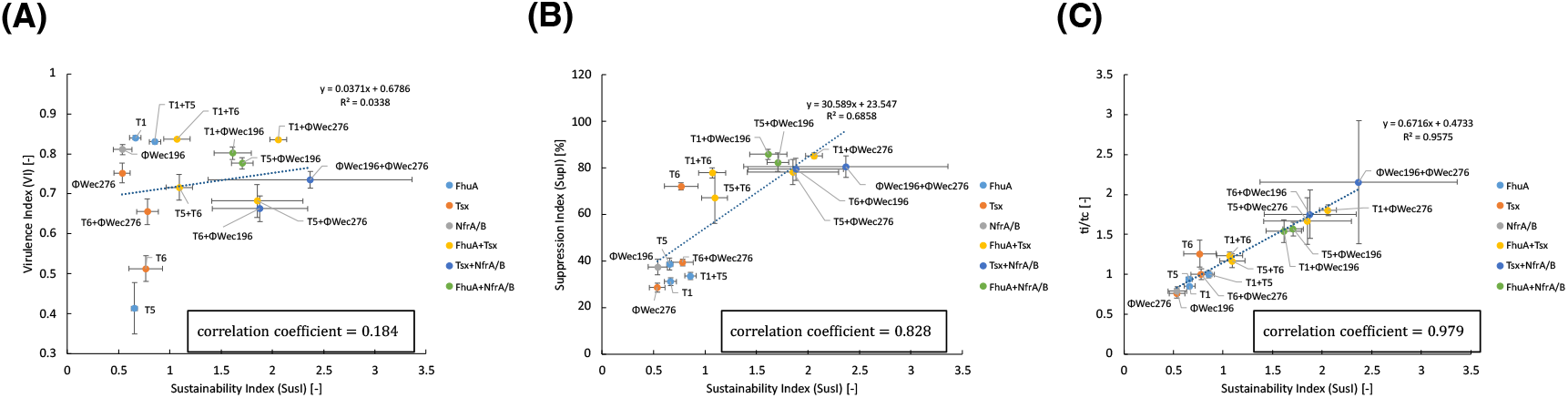
Correlation analysis between SusI and existing indices for phages infecting *E. coli* MG1655. SusI values at MOI = 1, calculated from the mean turbidity curve across three biological replicates (corresponding to the black diamond (◆) values in Fig. 2), are plotted on the x-axis against VI (A), SupI at MOI = 1 (B), and *t_i_*/*t_c_* at MOI = 1 (C) on the y-axis; error bars represent standard error across three biological replicates, as in Fig. 2. Data points are color-coded by receptor target combination (light blue: FhuA-targeting; red: Tsx-targeting; gray: NfrA/B-targeting; yellow: FhuA + Tsx-targeting; blue: Tsx + NfrA/B-targeting; green: FhuA + NfrA/B-targeting). The dashed line represents the least-squares linear regression fit; the boxed value indicates the correlation coefficient (r).

### Phage isolation, evaluation, and inter-index correlation analysis for *E. coli* ESBL1064

When testing whether five phages (T1, T5, T6, ΦWec196, and ΦWec276) capable of infecting MG1655 could infect the clinical isolate ESBL1064, only T6 exhibited practical lytic activity in liquid co-culture (T1 and T5 showed no activity; ΦWec196 and ΦWec276 showed limited activity). Three additional phages infecting ESBL1064 were obtained: ΦWec430 was selected from our laboratory’s phage library, while ΦWec461 through ΦWec467 were newly isolated from urban sewage. Among those isolated phages, ΦWec461 and ΦWec464 were selected for further analysis based on their lytic activity. ΦWec462, ΦWec463, and ΦWec465 were excluded from index evaluation due to the absence of lytic activity in liquid co-culture (Fig. S3); ΦWec466 and ΦWec467, though isolated and purified, were not subjected to growth curve analysis and were therefore also excluded from index evaluation.

Receptor group estimation was performed for all ESBL1064-infecting phages identified or isolated in this study (T6, ΦWec430, ΦWec461, ΦWec462, ΦWec463, ΦWec464, ΦWec465, ΦWec466, and ΦWec467), including those excluded from index evaluation (see Fig. S4 for full results). Resistant mutants were generated for each phage other than the reference phage T6. All resistant strains were first screened by spot assay against all phages, and those that retained susceptibility to all phages were excluded as likely transient-resistance isolates. Adsorption assays were then performed on the remaining strains, and those confirmed to carry receptor mutations were selected for subsequent infection range analysis. Infection range assays of the selected receptor mutant strains showed that ΦWec461 and ΦWec464 could infect ΦWec430-resistant receptor mutant strains, and conversely, ΦWec430 could infect receptor mutant strains resistant to ΦWec461 or ΦWec464 (Fig. S4A). Furthermore, host range assays using the *E. coli* panel strain collection revealed that the host range pattern of ΦWec430 was clearly distinct from those of ΦWec461 and ΦWec464, further supporting the assignment of different receptor groups (Fig. S4B). Based on these results, phages infecting ESBL1064 were classified into at least three distinct receptor groups: the Tsx-family group containing T6 (known), the group containing ΦWec430, and the group containing ΦWec461 and ΦWec464.

Growth curve measurements and index calculations were performed under the same conditions as for MG1655 for the four single phages (T6, ΦWec430, ΦWec461, and ΦWec464) and six two-phage cocktails (Figs. S5 and S6). Single-phage SusI values at MOI = 1 were low for all phages tested: ΦWec461 (0.291 ± 0.032), ΦWec464 (0.179 ± 0.054), ΦWec430 (0.101 ± 0.028), and T6 (0.351 ± 0.008). Among cocktails, all except ΦWec461+ΦWec464 (0.280 ± 0.022) showed higher SusI than their constituent single phages, with T6+ΦWec430 (1.535 ± 0.175) and T6+ΦWec461 (2.776 ± 0.790) showing a markedly elevated value. VI was notably low for T6 (0.133 ± 0.072), intermediate for ΦWec430 (0.501 ± 0.049), and high for ΦWec461 (0.832 ± 0.003) and ΦWec464 (0.807 ± 0.003). SupI was low (18–26%) for all single phages, with substantial increases observed for cocktails containing T6 (T6+ΦWec461: 76.3 ± 6.3%; T6+ΦWec464: 80.6 ± 2.1%). As observed with MG1655, *t_i_*/*t_c_* was highest for cocktails containing T6, with T6+ΦWec461 showing the maximum value (2.064 ± 0.471).

The correlation coefficient between SusI and VI was 0.364, confirming their independence as observed with MG1655 (Fig. S7A). The correlation coefficient between SusI and SupI was 0.934, indicating moderate-to-strong correlation (Fig. S7B); however, conditions containing T6 in cocktail and conditions containing ΦWec430 in cocktail showed comparable SupI values (60–80%), while SusI clearly differed between these groups (T6+ΦWec461: 2.776 vs. ΦWec430+ΦWec461: 1.272). The correlation coefficient between SusI and *t_i_*/*t_c_* was 0.989, indicating very strong correlation (Fig. S7C); unlike MG1655, no cases were observed in which turbidity remained elevated during the lysis period.

### *In vivo* validation using a mouse systemic infection model

Prior to *in vivo* validation, the infection and treatment model was established. *In vitro* antimicrobial susceptibility testing against ESBL1064 demonstrated resistance to β-lactam antibiotics (ampicillin, amoxicillin, and cefalexin), with effective sustained growth suppression achieved only by the carbapenem meropenem (Fig. S8A). Intraperitoneal administration of ESBL1064 to female BALB/c AJcl mice (6 weeks old) at 1 × 10^9^ CFU/mouse (100 µL) caused 100% mortality on day 1, establishing this dose as the infection model (Fig. S8B). To determine an effective antibiotic dose, meropenem was administered by tail vein injection at 10, 20, or 30 mg/kg (in 100 µL PBS) simultaneously with infection; only the 30 mg/kg dose resulted in 100% 7-day survival, and this dose was therefore designated as the positive control (PC) for subsequent experiments (Fig. S8C).

Following model establishment, three phage conditions with distinct SusI and VI profiles were selected for *in vivo* efficacy comparison: T6 alone (VI = 0.133, SusI = 0.351 at MOI = 1; low VI, low SusI), ΦWec461 alone (VI = 0.832, SusI = 0.291 at MOI = 1; high VI, low SusI), and the T6+ΦWec461 cocktail (VI = 0.818, SusI = 2.776 at MOI = 1; high VI, high SusI). Each phage preparation was administered simultaneously with infection at 1×10^9^ PFU/mouse (MOI = 1, 100 µL, tail vein injection) (n = 5). Both T6 alone and ΦWec461 alone resulted in 100% mortality within 2 days post-administration, whereas all 5/5 mice in the T6+ΦWec461 group survived for 7 days, matching the PC group (Fig. 4A).

**Fig. 4.**
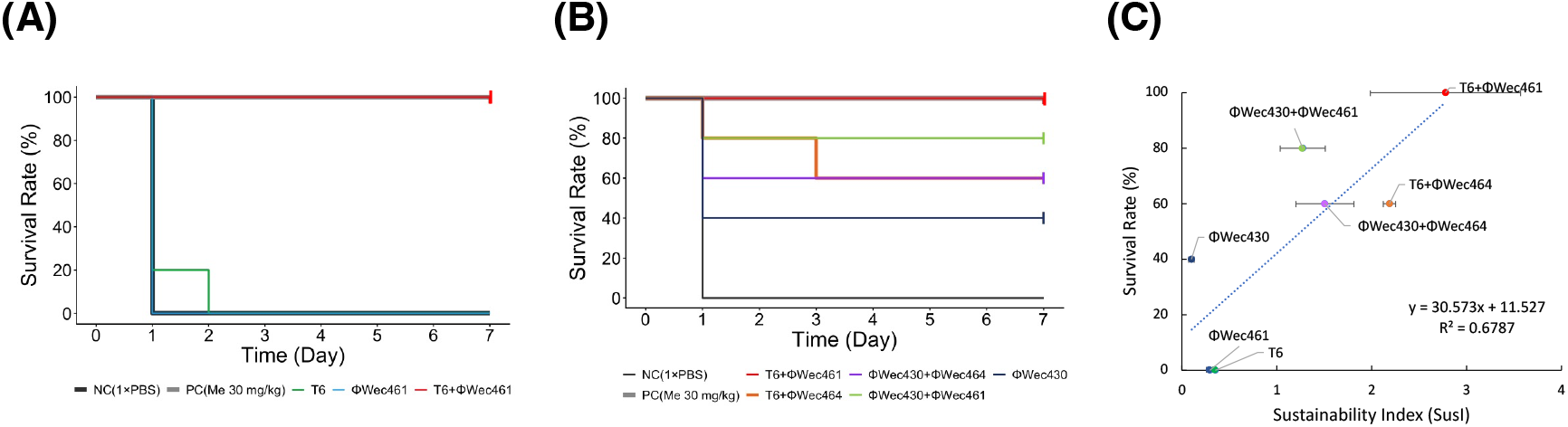
*In vivo* therapeutic efficacy of phage cocktails in a mouse model of systemic infection. (A) Kaplan–Meier survival curves of ESBL1064-infected mice following administration of three conditions with distinct SusI and VI profiles: T6 alone (low VI, low SusI), ΦWec461 alone (high VI, low SusI), and the T6+ΦWec461 cocktail (high VI, high SusI). The positive control (PC) group, treated with meropenem (30 mg/kg), is also shown. (B) Kaplan–Meier survival curves following administration of five conditions selected in descending order of SusI: four cocktails with consistently high VI (≈0.8), together with the constituent single phage with the lowest SusI (ΦWec430 alone), which also extended the range of VI examined. (C) Relationship between 7-day survival rate and SusI across both phage administration experiments. Each data point represents one condition. The solid line represents the regression line for the combined data from both experiments. n = 5 animals per group.

Subsequently, five conditions spanning a wide range of SusI values were selected to more systematically examine the relationship between SusI and survival: four cocktails (SusI = 1.272 ∼ 2.776) with consistently high VI (≈0.8) but differing SusI, together with the constituent single phage with the lowest SusI (ΦWec430 alone, VI = 0.501, SusI = 0.101), which also extended the range of VI examined. All were administered using the same protocol (n = 5). Day-7 survival was 5/5 for T6+ΦWec461, 3/5 for T6+ΦWec464, 3/5 for ΦWec430+ΦWec461, 2/5 for ΦWec430+ΦWec464, and 1/5 for ΦWec430 alone (Fig. 4B). Across both experiments, a positive relationship was observed between 7-day survival rate and SusI (Fig. 4C).

## DISCUSSION

In this study, we proposed SusI as a novel metric for quantitatively evaluating the sustainability of primary phage-mediated lysis and compared it with existing metrics (VI, SupI, and *t_i_*/*t_c_*) using the laboratory *E. coli* strain MG1655 and the clinical isolate ESBL1064. The results demonstrated that SusI captures information independent of VI, as VI primarily reflects bactericidal capacity during the initial phase of infection whereas SusI captures the persistence of lytic activity through to resistance emergence, and displays moderate-to-strong correlation with SupI and *t_i_*/*t_c_* while complementing aspects that the existing metric misses. Furthermore, phage cocktails with higher SusI values showed superior therapeutic efficacy in a mouse systemic infection model using ESBL1064, suggesting that SusI may be useful for predicting *in vivo* therapeutic outcomes.

### SusI captures information independent of VI

In both MG1655 and ESBL1064, the correlation coefficient between SusI and VI was approximately 0.2, confirming that the two metrics are independent. Previous studies have reported a high correlation (r = 0.94) between VI and PhageScore (18), indicating that these metrics assess similar aspects of the acute phase of phage–bacterium interaction. The independence of SusI from VI in the present study indicates that SusI captures distinct biological events that are not merely extensions of these existing metrics. This independence is explained by the difference in their integration intervals. VI references the entire growth curve from phage addition through *t_c_*, primarily reflecting rapid bacterial reduction during early infection, a process in which the phage adsorption rate constant for the target receptor is likely to play a major role, although this parameter was not directly measured in the present study and remains an important target for future investigation. SusI, by contrast, integrates only the primary lysis period from *t*_0_ to *t_i_*, capturing not just the absolute bactericidal capacity during early infection but a fundamentally different dimension: the persistence of lytic activity until resistance emergence. This reflects not only the phage’s intrinsic killing kinetics but also the population-level dynamics of bacterial resistance evolution, a process which unfolds after VI’s measurement window has already ended. Indeed, T1 and ΦWec196, despite having high VI values (0.840 and 0.811, respectively), showed relatively low SusI as single phages (0.664 and 0.539 at MOI = 1, respectively), demonstrating that superior early bactericidal activity does not necessarily confer sustained primary lysis. By combining these two metrics, it becomes possible to distinguish phages with strong adsorption capacity but limited lytic persistence from those with sustained lytic activity, enabling a more comprehensive characterization of phage lytic potential.

### SusI shows moderate correlation with SupI but provides unique information

The correlation coefficients between SusI and SupI were 0.828 for MG1655 and 0.920 for ESBL1064, indicating that the two metrics are related but not equivalent. SupI has been defined as the percentage of growth suppression over a fixed 30-hour period (20) and has been widely used to capture the overall antibacterial efficacy of phages. While acknowledging its utility, the present study identified two scenarios in which SupI falls short of what SusI can capture. First, there is the problem of overestimation due to delayed lysis initiation. T6 alone showed a high SupI (≈72%) but a relatively low SusI (≈0.77) against MG1655 among single phages. Growth curve analysis revealed that T6 exhibits delayed lysis initiation compared to other phages, with a smaller difference from the control during the early measurement period. Because SupI integrates the area difference over the entire measurement period, it yields a high value by capturing the strong suppression achieved during the later phase, despite the early lag. SusI evaluates only the area difference from *t*_0_ onwards and therefore avoids this confound. Since rapid action is an important clinical property of phage therapy, the possibility that SupI may not accurately reflect the speed of lysis onset is a relevant consideration in cocktail design. Second, SupI can be confounded by growth dynamics occurring outside the primary lysis period. Secondary lysis (a re-decline in turbidity following resistance emergence) was observed for T1+T6. The SupI of ≈78% for this condition includes the contribution of this secondary lysis event, whereas SusI of ≈1.07 integrates only the primary lysis period from *t*_0_ to *t_i_*, excluding secondary lysis. The phenomenon in which bacteria acquiring resistance to one phage are subsequently lysed by the other is biologically interesting as a feature of cocktails combining phages targeting different receptors. However, when the goal is to compare the primary lytic sustainability of different cocktails for rational design purposes, SusI is a more appropriate metric than SupI, which can be confounded by secondary lysis.

### Strong correlation between SusI and *t_i_*/*t_c_*, and their differences

The correlation coefficients between SusI and *t_i_*/*t_c_* were 0.979 for MG1655 and 0.989 for ESBL1064, indicating that the two metrics largely capture similar information. This is likely because the numerator of SusI tends to be dominated by the time span from *t*_0_ to *t_i_*, suggesting that lysis duration is the primary determinant of SusI. However, there are cases where SusI provides information beyond *t_i_*/*t_c_*. T6 alone with MG1655 is a prime example: despite a high *t_i_*/*t_c_* (1.258), SusI (0.766) fell below the regression line. Growth curve analysis confirmed that T6 maintained a higher turbidity level during the lysis period compared to other phages, reflecting weaker lytic activity per unit time. Because *t_i_*/*t_c_* considers only the duration of lysis, it can yield high values even when lytic intensity is weak but prolonged. SusI, by additionally incorporating the area difference between phage-treated and control groups during the lysis period, integrates both time and lytic intensity and is thus conceptually superior in this regard.

By contrast, no discrepancy between *t_i_*/*t_c_* and SusI was observed among the ESBL1064 phages. This may reflect phenotypic bias arising from the fact that phages infecting ESBL1064 were isolated from a single host using the double-layer agar plaque method. Phages isolated from a single host are known to converge on genetically similar clones capable of evading that host’s defense mechanisms (27, 28), potentially limiting the phenotypic diversity of lytic activity in the resulting collection. When phages with more diverse lytic characteristics are used (e.g., those obtained through alternative isolation methods such as liquid co-culture or from different host strains), more pronounced discrepancies between SusI and *t_i_*/*t_c_* would be expected to emerge.

### SusI elevation through different-receptor cocktails and the mechanism of resistance suppression

In both MG1655 and ESBL1064, cocktails combining phages targeting different receptors consistently showed higher SusI than same-receptor cocktails or single phages. This can be interpreted as follows: because phages targeting different receptors are present simultaneously, bacterial strains require multiple receptor mutations at the same time to acquire resistance, thereby delaying resistance emergence and extending *t_i_*; in other words, the selective pressure toward resistance becomes multifaceted. This interpretation is consistent with previous studies demonstrating reduced resistance emergence rates with different-receptor phage cocktails (29–31). The observation that T1+T5, a same-receptor cocktail, showed SusI values comparable to those of its constituent single phages further supports the importance of receptor diversity. For ESBL1064 phages, cross-infection assays of resistant strains (including confirmed receptor mutant strains) and host range analyses using our *Enterobacteriaceae* panel strain collection independently confirmed that ΦWec430 and ΦWec461/ΦWec464 belong to distinct receptor groups, each also distinct from the Tsx receptor targeted by T6. The high SusI values of T6+ΦWec461 and T6+ΦWec464 are consistent with these being combinations targeting different receptors. On the other hand, ΦWec430+ΦWec461 is also a different-receptor combination, yet showed lower SusI than cocktails containing T6, and this pattern held across all ΦWec430-containing cocktails (T6+ΦWec430: 1.535; ΦWec430+ΦWec461: 1.272; ΦWec430+ΦWec464: 1.504), each falling below T6+ΦWec461 (2.776) and T6+ΦWec464 (2.187) despite comparable receptor diversity. This ceiling effect parallels the comparatively low single-phage SusI of ΦWec430 (0.101 ± 0.028) relative to the other three phages (0.179 – 0.351), rather than its VI, which was in fact intermediate (0.501 ± 0.049) and not the lowest among the four phages. This suggests that receptor diversity alone is not sufficient to elevate cocktail SusI: the constituent phages must also individually sustain effective lytic pressure once co-administered, and a phage with intrinsically brief primary lysis, such as ΦWec430, may limit the achievable SusI of the resulting cocktail even when paired with a phage targeting a distinct receptor (19, 25). This case illustrates how SusI can complement receptor diversity information by capturing aspects of cocktail efficacy that receptor classification alone cannot predict. It should be noted that T6 infection of ESBL1064 was limited to partial lysis in liquid culture without plaque formation, suggesting that the efficiency of T6 infection through the Tsx receptor on ESBL1064 is lower than that on MG1655. This may partly explain why T6 alone showed low VI (0.133) and SusI (0.351) against ESBL1064, with high SusI emerging only upon cocktail formation.

The utility of combining phages targeting different receptors to suppress resistance emergence is well established, and methods for efficiently selecting such phages have been reported. However, the present study using ESBL1064 demonstrated that even among different-receptor combinations, clear differences exist in both cocktail SusI and *in vivo* therapeutic efficacy. Despite both being different-receptor combinations, T6+ΦWec461 (SusI ≈ 2.78) substantially outperformed ΦWec430+ΦWec461 (SusI ≈ 1.27), with a corresponding difference observed *in vivo*. Previous studies have suggested that incorporating phage-intrinsic physiological properties such as infection range or lysis kinetics can facilitate the design of effective cocktails (25, 26). Among such properties, SusI uniquely integrates the persistence and depth of lysis in a single quantitative measure. By adding the dimension of lytic quality to the premise of receptor diversity, SusI is expected to serve as a complementary metric enabling more precise design of phage cocktails effective for clinical applications.

### Predictive validity of SusI in a mouse systemic infection model

Whether SusI evaluated *in vitro* can predict *in vivo* therapeutic efficacy is a fundamental question for the practical implementation of phage therapy. In this study, only T6+ΦWec461 (the cocktail with the highest SusI) achieved 100% 7-day survival, while T6 alone and ΦWec461 alone showed no therapeutic efficacy. These results suggest that *in vitro* SusI may reflect *in vivo* therapeutic outcome. It is particularly notable that, despite markedly different VI values between T6 (VI = 0.133) and ΦWec461 (VI = 0.832), neither was effective as a single phage while their cocktail achieved 100% survival, demonstrating that early bactericidal capacity as measured by VI alone cannot predict therapeutic success. Furthermore, the complete 7-day survival achieved with T6+ΦWec461 was reproduced in two independent experiments (5/5 mice in each), supporting the robustness of this finding.

In the experiment comparing the five highest-SusI cocktails, a general positive trend was observed between 7-day survival rate and SusI, although complete rank correspondence was not achieved. One possible explanation for this discrepancy is that phage-intrinsic properties of ΦWec461 (such as genome size or interactions with host immunity) contributed to survival outcomes independently of SusI. The tendency for higher survival rates in cocktails containing ΦWec461 is consistent with this possibility. It is well established that phage efficacy in animal models depends on multiple factors that are difficult to evaluate *in vitro*, including susceptibility to clearance by the mammalian immune system and resistance to bacterial defense mechanisms, which may have contributed to the rank discrepancy. Whole-genome sequencing and receptor identification of ΦWec461 in particular represent important steps toward characterizing such phage-specific *in vivo* properties, including genes relevant to immune evasion and circulatory stability. Although the limitation of a small sample number (*n* = 5) cannot be excluded as a contributing factor, the consistent positive relationship between SusI and survival observed across both experiments provides preliminary evidence that SusI can function as a predictive indicator of *in vivo* therapeutic efficacy.

### Limitations and future perspectives

The results presented here suggest that SusI can serve as a consistently useful metric spanning the continuum from *in vitro* phage evaluation to prediction of *in vivo* therapeutic efficacy. However, several important limitations warrant consideration. First, the very high correlation between SusI and *t_i_*/*t_c_* indicates that the conditions under which SusI provides clearly additive information over *t_i_*/*t_c_* may be limited to phage panels with sufficient phenotypic diversity in lytic activity. Future work should quantitatively evaluate the relationship between phage physiological parameters (such as adsorption rate constant and burst size) and SusI, to more precisely define the unique information that SusI captures. Second, the target receptors and complete genome sequences of ΦWec430, ΦWec461, and ΦWec464 have not yet been identified. Consequently, the mechanistic interpretation of why receptor diversity elevates cocktail SusI against ESBL1064 remains an inference based on findings from MG1655 phages, and verification through future genome sequencing and receptor identification experiments is warranted. Third, the mouse infection model in this study was limited to preliminary experiments with *n* = 5. Establishing the predictive validity of SusI will require large-scale *in vivo* studies incorporating a broader range of clinical isolates and phage collections. Despite these limitations, the present study is significant as the first demonstration that SusI, a metric calculable from in vitro growth curve data, can correspond to in vivo therapeutic outcomes. In the development of novel antibacterial treatments, simple metrics capable of predicting therapeutic efficacy at the screening stage are greatly needed, and SusI represents a strong candidate for this role. As research toward the clinical implementation of phage therapy continues, SusI is expected to serve as a practical bridge between in vitro screening and in vivo efficacy evaluation.

## Acknowledgements

We thank the staff at Gunma University Hospital for providing clinical isolates including ESBL1064. We are grateful to Akari Chiba (Waseda University) for isolating and providing ΦWec430. We thank Han Guangli, Shouta Katayama, and Yuki Takei for their invaluable assistance with mouse tail vein injection experiments. We are grateful to Dr. Kazuki Kitaoka for his advice on antibiotic selection and dose determination for the mouse experiments.

This work was supported by the Waseda University Grant for Special Research Projects (project number: 2025C-509), the Takahashi Industrial and Economic Research Foundation (fiscal years 2025 and 2026), the Waseda University Faculty of Science and Engineering Research Institute Young Researcher Support Program (Early Bird Program), a Grant-in-Aid for Scientific Research (B) from the Japan Society for the Promotion of Science (JSPS; grant number: 23K27412), and a JSPS Grant-in-Aid for Early-Career Scientists (grant number: 26KJ0420).

Claude (Anthropic) and Gemini (Google) were used to assist with English language editing. Use of these tools was limited to refining written expression; the accuracy of all scientific content in this manuscript was verified by the authors.

